## Supplementary material for "An *in vitro* human vessel model to study *Neisseria meningitidis* colonization and vascular damages": Supp info

### Supplementary Figures

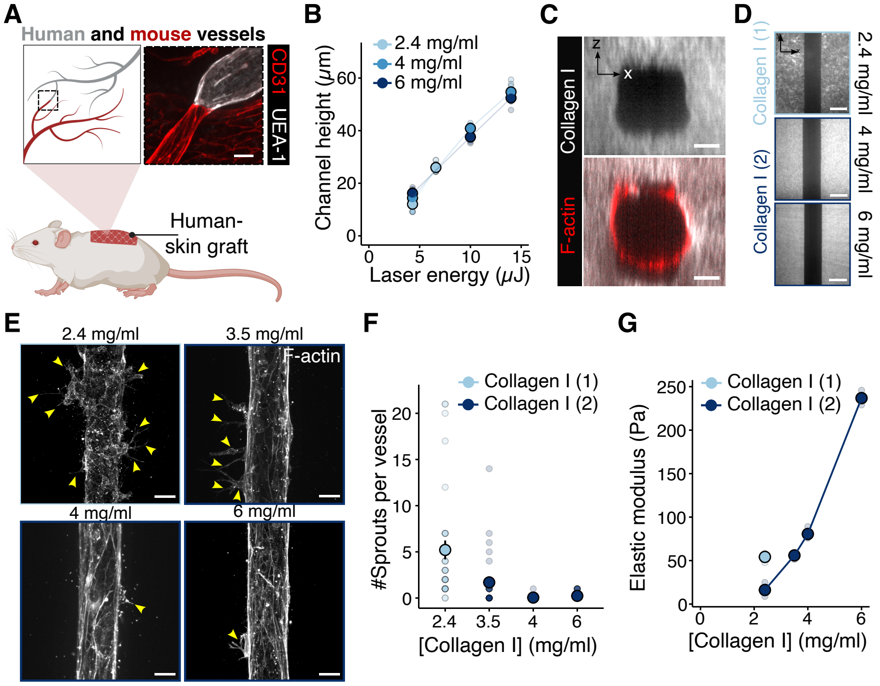


**Supplementary Figure 1. Development, optimization, and characterization of a tissue-engineered Vessel-on-Chip platform based on photoablation**. (A) Schematic representation of the *in vivo* conditions assessed in this study: human vessels imaged in the human-skin xenograft. Scale bar: 30 µm. (B) Channel height depends on UV-Laser energy for 2.4 mg/ml (FujiFilm), 4 mg/ml (Corning), and 6 mg/ml (Corning) collagen I gels. (C) Orthogonal view of a bear and endothelial-layered collagen-carved tube. Scale bar: 20 µm. (D) Collagen-carved scaffold for 2.4 mg/ml (FujiFilm), 4 mg/ml (Corning), and 6 mg/ml (Corning) collagen I gel before cell seeding. Scale bar: 50 µm. (E) Confocal image of the VoC in 2.4 mg/ml - 3.5 mg/ml - 4 mg/ml and 6 mg/ml collagen I. Fujifilm collagen I has been used to make the 2.4 mg/ml (cyan circle) and Corning collagen I has been used to make the solution from 3.5 mg/ml to 6 mg/ml (blue circle). Scale bar: 30 µm. (F) Graph representing the number of sprouts per vessel for four collagen I concentrations: 2.4 mg/ml (FujiFilm), 3.5 mg/ml, 4 mg/ml, and 6 mg/ml (Corning). Each dot represents one vessel. For each condition, the mean ± s.d. is represented (2.4 mg/ml: 5.20 ± 6.0 (n=28) — 3.5 mg/ml: 1.69 ± 3.04 (n=29) — 4 mg/ml: 0.045 ± 0.22 (n=21) — 6 mg/ml: 0.23 ± 0.43 (n=30)) (G) Visco-elasticity measurements of collagen gels with a rheometer. Each dot represents a measure of each gel. For each condition, the mean ± s.d. is represented (FujiFilm, 2.4 mg/ml: (n=3) — Corning, 2.4 mg/ml: (n=3) - 3.5 mg/ml: (n=3) - 4 mg/ml: (n=3) - 6 mg/ml: (n=3)). The mentions (1) and (2) in (D), (F), and (G) refer to the providers of the collagen I, FujiFilm and Corning, respectively.

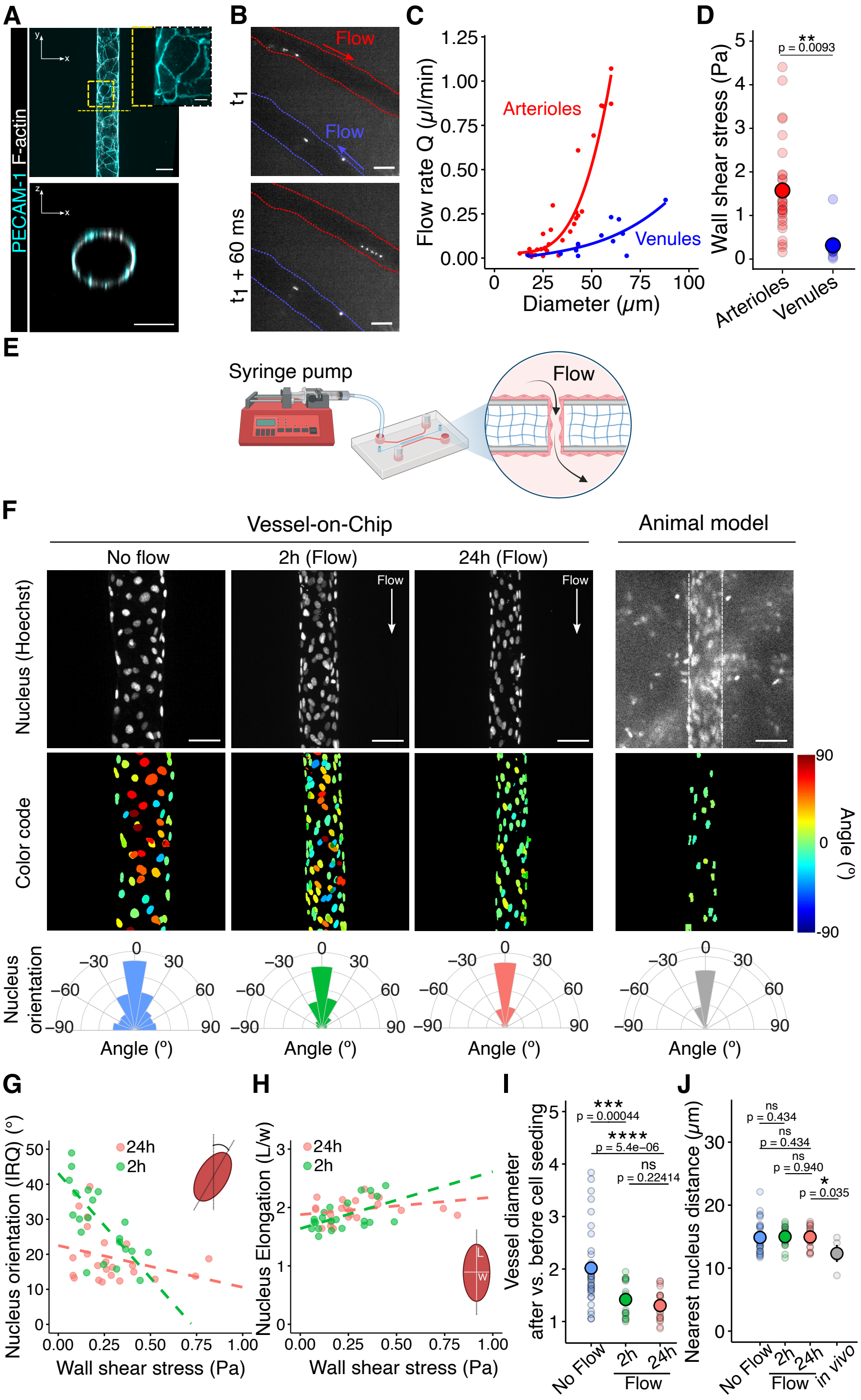

**Supplementary Figure 2. Endothelial cells in the Vessel-on-Chip devices respond to flow-induced shear stress**. (A) Confocal images of PECAM-1 staining of the VoC. Scale bar: 50 µm. (B) Intravital images of PIV tracking of an arteriole and a venule in the mouse model (ear skin dermis). Scale bar: 50 µm. (C) Graph representing the flow rate depending on the vessel size. The red and blue fitting curves follow the $y = a/x^3$ equation for arterioles and venules, respectively. (D) *In vivo* wall shear stress of blood flow in arterioles and venules in the mouse model. Each dot represents a vessel. For each condition, the mean ± s.d. is represented (Arterioles: 1.57 ± 1.05 Pa (n=31) – Venules: 0.31 ± 0.35 Pa (n=12)). (E) Schematic of VoC experiments under flow conditions, controlled with a syringe pump. (F) Representative images of nucleus alignment (top) with the corresponding segmented/color-coded image (middle) and the circular plot of orientation (bottom). Scale bar: 30 µm. (G) Graph of nucleus interquartile range (orientation) and (H) nucleus elongation according to WSS. (I) Vessel diameter ratio before and after cell seeding ($D_{after}/D_{before}$) and (J) nearest nucleus distance for no flow condition, after 2h and 24h of flow. Each dot corresponds to a vessel. For each condition, the mean ± s.d. is represented (No flow conditions: 2.02 ± 0.74 and 14.90 ± 2.40 µm (n=34) – 2h of flow: 1.42 ± 0.31 and 15.00 ± 1.52 µm (n=22) – 24h of flow: 1.30 ± 0.27 and 14.9 ± 1.55 µm (n=26) — Animal model: 12.30 ± 2.70 µm (n=4)). All statistics have been computed with Wilcoxon tests.

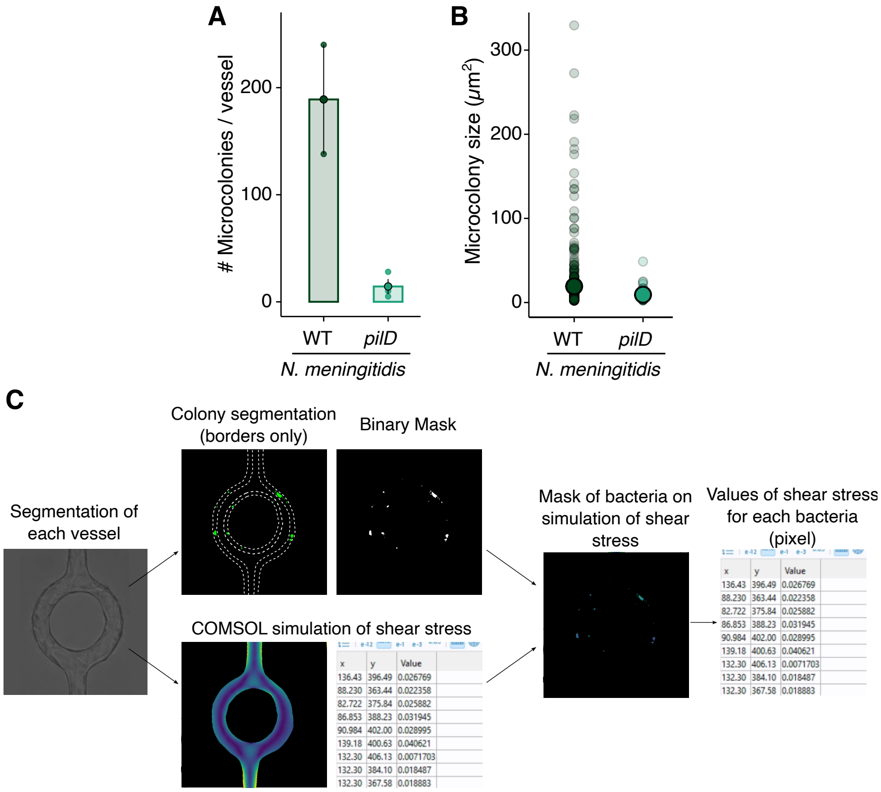


**Supplementary Figure 3.** ***Neisseria meningitidis* use T4P to adhere on the Vessel-on-Chip and its adhesion does not depend on the difference of shear stress.** (A-B) Graphs representing the numbers and the sizes of microcolonies within the VoC infected with WT or *pilD Nm* strains. For each condition, the mean ± s.d. is represented (WT: 189.0 ± 72.1 (n=2 vessels) and 16.60 ± 3.39 μm$^2$ (n=378 microcolonies) – *pilD*: 14.3 ± 12.1 (n=3 vessels) and 9.03 ± 4.47 μm$^2$ (n=43 microcolonies)). (C) Analysis pipeline of the correlation between bacterial location and shear stress in Vessel-on-Chip.

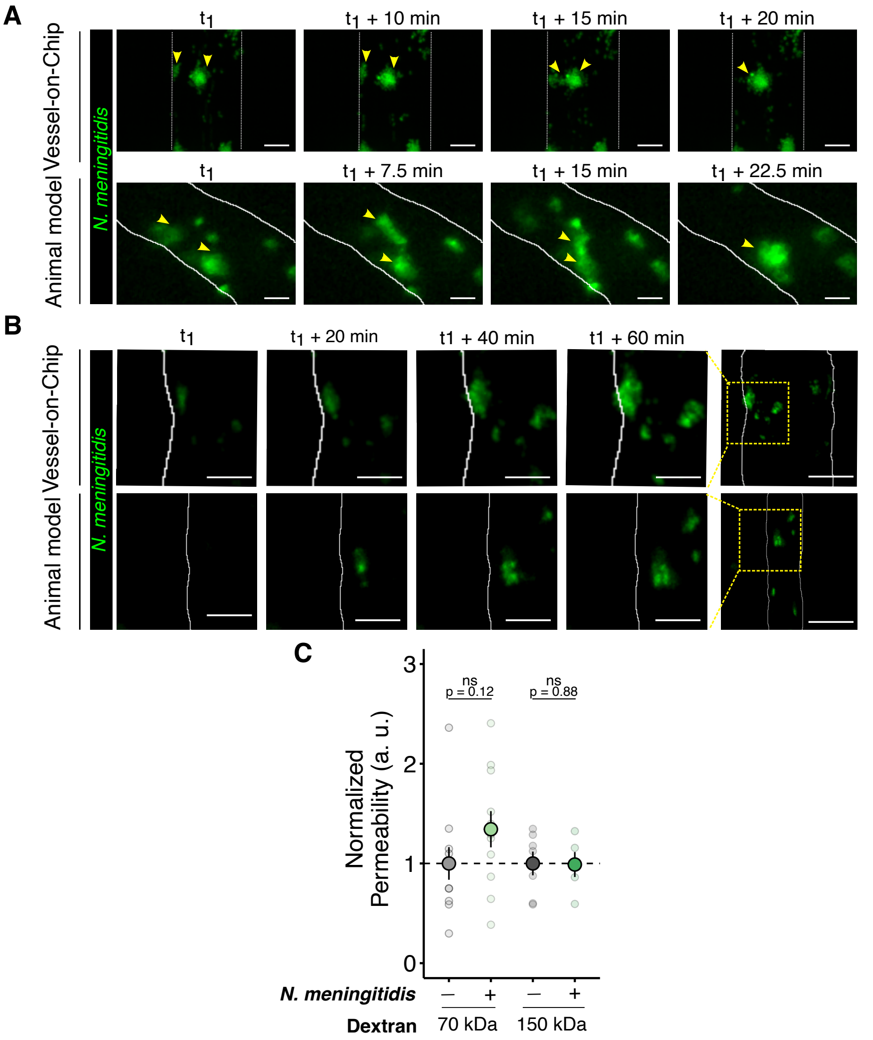


**Supplementary Figure 4.** ***N. meningitidis*** **colonies grow and reorganize** ***in vitro*** **under flow conditions, and** ***in vivo*** **permeability is not affected by** ***N. meningitidis*** **at early stages of infection**. (A) Time-lapse images of bacterial growth in a VoC and an infected human vessel in the skin-xenografted mouse model. Scale bar: 30 µm. (B) Zoom on a bacterial fusion event in a VoC (top) and in a human skin-grafted mouse model (bottom). Scale bar: 10 µm. (C) Permeability of infected human vessels in the human skin-xenografted mouse model (green), normalized with the permeability of non-infected vessels (gray). For each condition, the mean ± s.d. is represented (70 kDa - Control: 1 ± 0.542 (n=11 vessels, N=3 mice), *Nm*: 1.34 ± 0.603 (n=11 vessels, N=3 mice) — 150 kDa - Control: 1 ± 0.314 (n=7 vessels, N=3 mice), *Nm*: 0.989 ± 0.280 (n=5 vessels, N=3 mice). Statistics have been done with Wilcoxon tests.

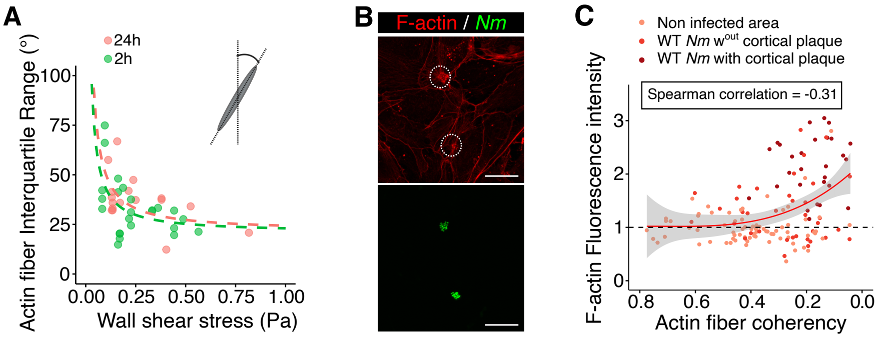


**Supplementary Figure 5. Flow-induced aligned actin stress fibers are reorganized below** ***N. meningitidis*** **microcolonies**. (A) Interquartile range of actin fibers according to the flow-induced wall shear stress. (B) Confocal images of F-actin cortical plaque at the infection sites in the 2D regions of the chips (n=5 chips, n=3 experiments). Scale bar = 40 µm. (C) Graph representing the correlation between F-actin mean fluorescence intensity and actin fiber coherency.

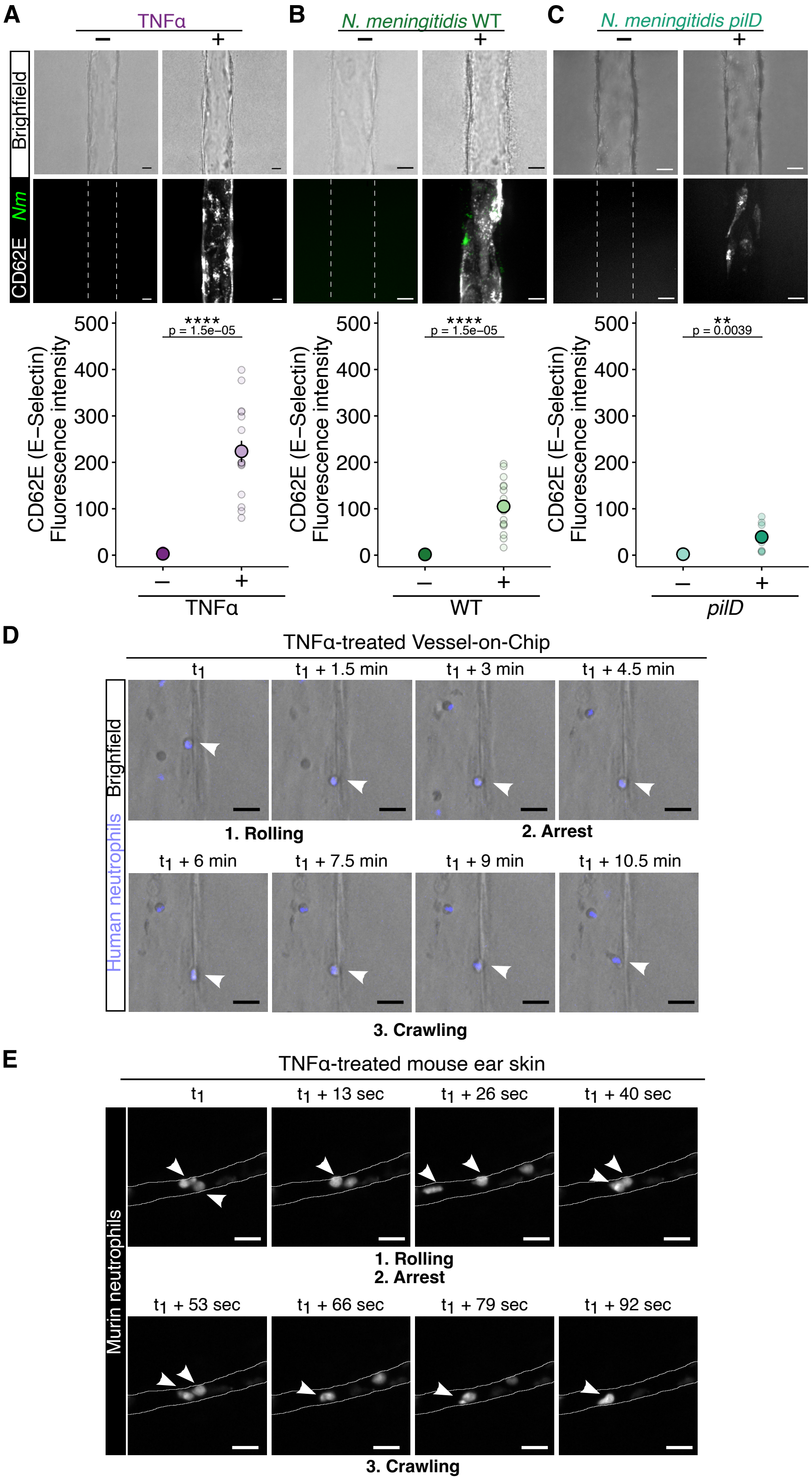


**Supplementary Figure 6. The infected Vessel-on-Chip model recapitulates the human neutrophil response to *N. meningitidis* infection**. (A-C) Brightfield and fluorescence confocal images of a VoC before (-) and after (+) 4h of TNF$\alpha$ treatment, or infection of WT or *pilD Nm* strains (top). Graph of the E-selectin mean fluorescence intensity quantification for each condition (bottom). Each dot represents a vessel. For each condition, the mean ± s.d. is represented (TNF$\alpha$ - Before treatment: 1.85 ± 3.85 – 4h post treatment: 224.0 ± 93.7 (n=17) — WT - Before infection: 1.42 ± 1.02 – 4h p.i.: 105.0 ± 54.4 (n=17) — *pilD* - Before infection: 1.42 ± 1.02 – 4h p.i.: 105.0 ± 54.4 (n=9)) Statistics have been computed with t-tests. (D) Time-lapse of the adhesion cascade of human neutrophils on the human endothelium (VoC). (E) Time-lapse of the adhesion cascade of mouse neutrophils on the mouse endothelium (mouse ear skin).
